# Control Theory Concepts for Modeling Uncertainty in Enzyme Kinetics of Biochemical Networks

**DOI:** 10.1101/618777

**Authors:** Ljubisa Miskovic, Milenko Tokic, Georgios Savoglidis, Vassily Hatzimanikatis

## Abstract

Analysis of the dynamic and steady-state properties of biochemical networks hinge on information about the parameters of enzyme kinetics. The lack of experimental data characterizing enzyme activities and kinetics along with the associated uncertainties impede the development of kinetic models, and researchers commonly use Monte Carlo sampling to explore the parameter space. However, the sampling of parameter spaces is a computationally expensive task for larger biochemical networks. To address this issue, we exploit the fact that reaction rates of biochemical reactions and network responses can be expressed as a function of displacements from thermodynamic equilibrium of elementary reaction steps and concentrations of free enzymes and their intermediary complexes. For a set of kinetic mechanisms ubiquitously found in biochemistry, we express kinetic responses of enzymes to changes in network metabolite concentrations through these quantities both analytically and schematically. The tailor-made sampling of these quantities allows for characterizing the missing kinetic parameters and accelerating the efforts towards building genome-scale kinetic metabolic models.

## 1. INTRODUCTION

Evolutionary processes in biological systems gave rise to a wide range of control and regulatory mechanisms to ensure their survival and robustness under varying environmental conditions. For a comprehensive understanding of cellular organisms, we have to consider them at the system-wide level and use the appropriate analytical methods.^*1*^ The methods from systems and control theory^*2*–*5*^ are particularly suitable for the analysis of biological systems, and many significant properties of cellular organisms have been discovered using these concepts.^*6, 7*^

The theoretical analysis and study of metabolic processes and ways to control fluxes in metabolic networks have been mostly focused on developing the quantitative descriptions of metabolism.^*8*^ A predominant approach in these studies is Metabolic Control Analysis (MCA), a parametric sensitivity analysis of a metabolic system around a steady-state.^*9–12*^ Though the theory of MCA is well developed, information about kinetic properties of enzymes that is required for its successful application is scarce.

To obtain the values of kinetic parameters, one can use experimental data and perform parameter estimation,^*13–16*^ or explore the parameter space by employing Monte Carlo sampling techniques,^*16, 17*^ The latter approach is prevalent in newer kinetic modeling methods.^*18*^ However, the random sampling of kinetic parameters spaces of large biochemical networks becomes computationally challenging as the size of network is growing.^*17*^ Efficient methods for exploring large parametric spaces of biochemical networks require tailor-made sampling techniques that exploit the specific structure of the networks while considering physico-chemical constraints.^*17*, *19*^ Recently, a novel method for characterization and reduction of uncertainty in kinetic models was proposed that further alleviates issues with intensive computational requirements by identifying parameters and their bounds relevant for the analyzed physiology.^*20*^

In our previous work^*19*^, we proposed a framework for modeling of uncertainty in the enzyme kinetics and efficient sampling technique that allows us to sample parametric space of large biochemical networks. The framework allows us to calculate the local parametric sensitivity coefficients of biochemical reactions, dubbed elasticities within the Metabolic Control Analysis (MCA) formalism^*9–12*^, as a function of the distribution of enzymes among their free form and intermediary complexes, the thermodynamic displacements of the reactions from the equilibrium, and the net reaction fluxes. The proposed formulation allowed us to explicitly integrate the conservation of the total amount of enzymes, metabolite concentrations and reaction thermodynamics, and perform an efficient Monte Carlo sampling for generating all states within modeled enzymatic mechanisms. The features of the proposed framework were illustrated through examples of the three-step reversible and irreversible Michaels-Menten kinetic mechanisms.

While the single-substrate single-product mechanisms can be used to model more complex enzymes, e.g., certain hydrolyses are commonly described with these mechanisms because water is abundant in living cells and its concentration is considered constant, according to a strict definition these mechanisms are rather infrequent in biochemistry as they are confined to isomerizations.^*21*^ In this work, we extend the previously proposed framework^*19*^ to more ubiquitous mechanisms appearing in biochemical networks such as ordered Bi Bi, ping pong Bi Bi, ordered Uni Bi and ordered Bi Uni mechanism. It is estimated that the Bi Bi mechanisms cover more than 60% of known enzymatic reactions.^*21*^ We also propose a novel schematic method for the derivation of the analytic expressions of elasticities allowing us to skip altogether algebraic manipulations that are particularly cumbersome for more complex mechanisms. The method is closely related to the signal flow graphs methods used in the analysis of (i) electronic circuits and control loops,^*22*^ and (ii) control and regulation in metabolic pathways.^*23–25*^ Furthermore, we present a bottom-up workflow that makes use of the computed elasticities to determine the steady-state outputs of biochemical networks induced by the changes in the enzyme activities despite uncertainties in kinetic properties of enzymes.

## 2. METHODS

### 2.1. (Log)Linear Description of Biochemical Networks and Metabolic Control Analysis

Consider a biochemical system with *n* metabolites involved in *m* enzymatic reactions. The mass balances of the system are described by

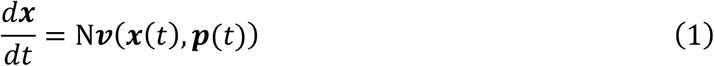

where the stoichiometric matrix, 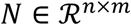, describes the network topology, 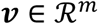 is the reaction rate vector, 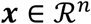 is the vector of metabolite concentrations, and 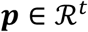 is the vector of manipulated inputs. Due to conservation relationships among metabolites in the network, the stoichiometric matrix is rank deficient, and the rows corresponding to dependent metabolites, 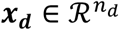, can be expressed as a function of the rows corresponding to independent metabolites, 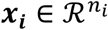, so that the system (Eq. 1) takes the form

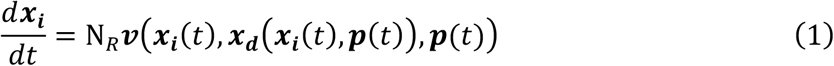

where N_*R*_ is the reduced stoichiometric matrix containing the rows corresponding to the independent metabolites.^*26*^

The vector of metabolic outputs, 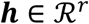, can be formed in general as a nonlinear function of reaction rates, **𝒗**, the independent metabolites, ***x***_***i***_, and the inputs, ***p***.^*8*, *12*^ For simplicity, here we choose ***h*** to contain the following two variables:

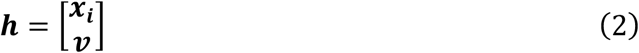

Following the derivation from Hatzimanikatis and Bailey^*12*^, we can linearize the system (Eqs. 1–2) around a steady state 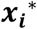, ***p***^*^ to obtain a (log)linear model:

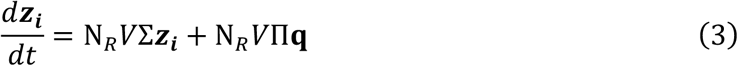

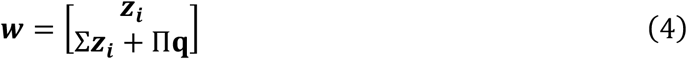

where the vectors ***z***_i_ and **q** represent the logarithmic deviations of the state variables and parameters whose elements are defined as:^*12*^

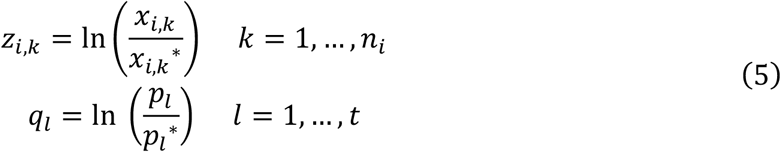

and the matrices *V*, Σ, and Π are:

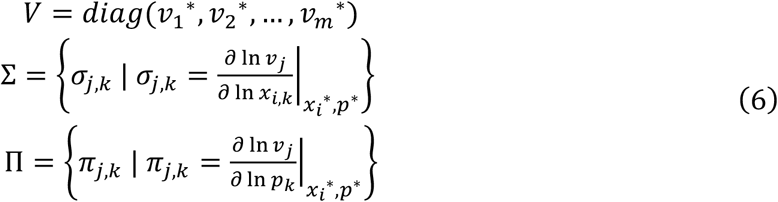

Hatzimanikatis and Bailey have demonstrated^*8, 12*^ that (log)linear models can accurately describe the dynamic behavior of metabolic responses. Moreover, they have shown that the flux and concentration control coefficients defined within the framework of Metabolic Control Analysis (MCA)^*9, 10, 12*^ can readily be derived from the (log)linear model (Eqs 3-6). Indeed, the flux and concentration control coefficients that quantify the responses of biochemical networks to the changes in systems parameters such as enzyme activities are the steady-state gains of the (log)linear system. By expressing the steady state solution of Eqs. 3-6, one obtains

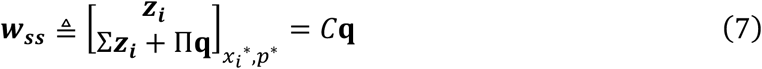

with

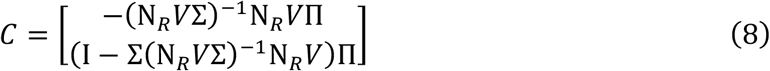

The matrix *C* is the control coefficient matrix of the outputs ***h*** (metabolite concentrations and metabolic fluxes) with respect to inputs ***p*** (within the MCA formalism^*9, 10, 12*^, the inputs ***p*** are considered as parameters as they do not change at the steady state). By inspecting Eq. 8, we observe that for computing the control coefficients we need to accurately determine the matrices of sensitivities of reaction rates with respect to the metabolite concentrations, Σ, and the parameters, Π (within the MCA formalism these matrices are called the elasticity matrices). The rates of reactions constituting a biochemical network are characterized by the mechanistic properties of catalyzing enzymes, but also by the states of the biochemical network, i.e., metabolic fluxes and metabolite concentrations.

Therefore, for determining the elasticity matrices Σ and Π, the steady-state values of fluxes and concentrations in the network are required. This task can be performed by employing methods that integrate information about thermodynamics in the context of Flux Balance Analysis^*27*^ such as the Thermodynamics-based Flux Analysis (TFA)^*28–30*^, the energy balance analysis (EBA)^*31*^, and the network-embedded thermodynamic analysis (NET analysis)^*32*^.

In contrast, determining the mechanistic properties of enzymes catalyzing the reactions involved in the networks is a challenging task because the comprehensive knowledge of the kinetic properties of enzymes is lacking.^*19, 33–35*^ To address this challenge, the space of enzyme states can be explored,^*19*^ and the information about the distribution of the enzymes among their free form and the intermediary complexes can then be used to compute the elements of the elasticity matrices Σ and Π. These matrices are, in turn, used to compute the control coefficients.

The set of procedures for determining the flux and concentration control coefficients is assembled in a conceptual workflow (Figure 1), and its constitutive elements are detailed below.

**Figure 1.**
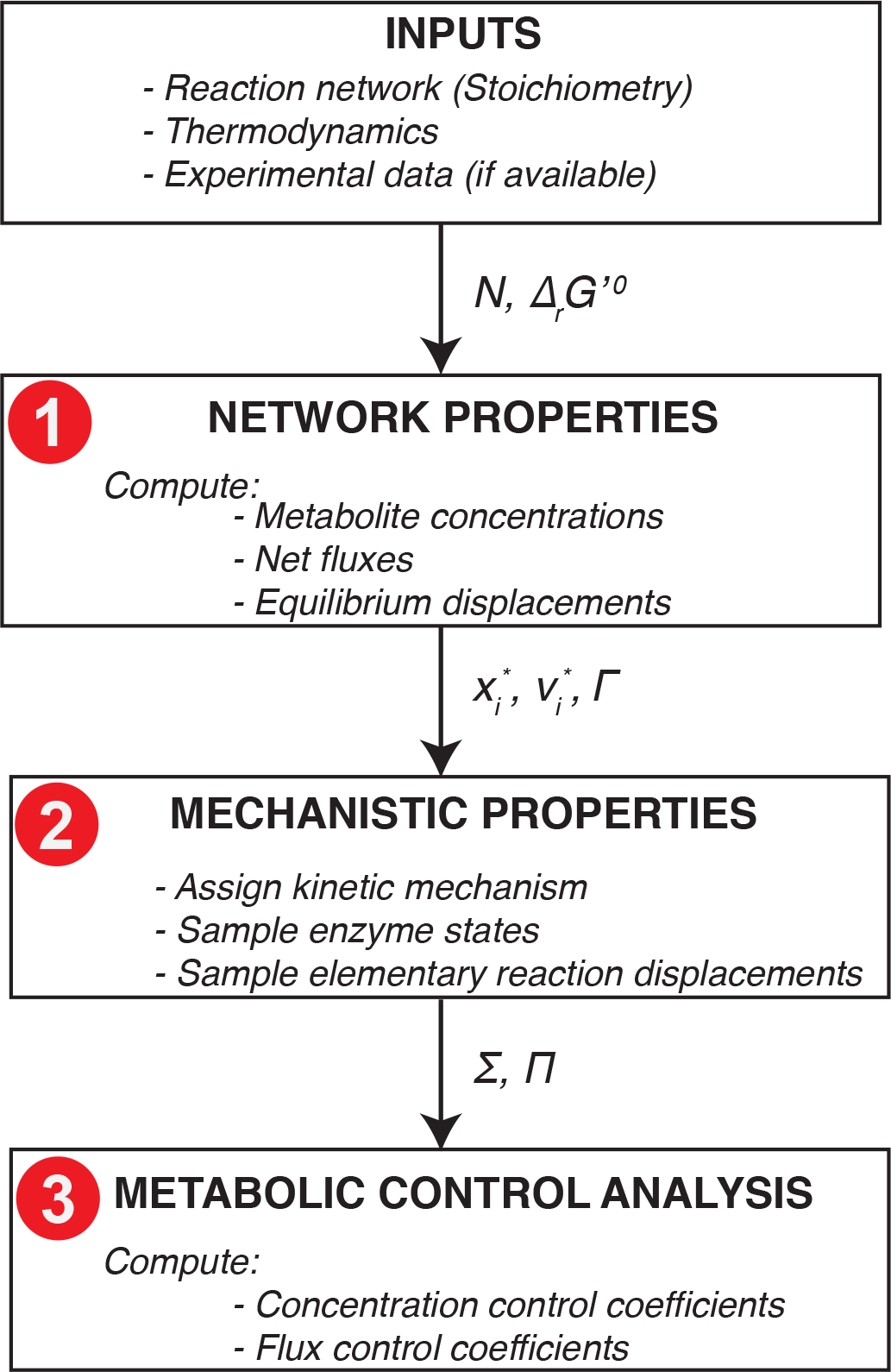
A workflow for the computation of flux and concentration control coefficients based on the Monte Carlo sampling of the enzyme states and elementary reaction displacements.

### 2.2. Displacements from Thermodynamic Equilibrium

The displacement of a reaction from its thermodynamic equilibrium is defined as follows:^*19, 36, 37*^

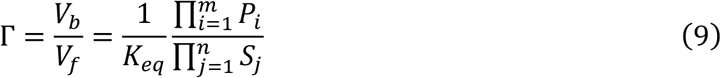

where *V*_*b*_ and *V*_*f*_ denote the backward and the forward reaction rates of the overall reaction, and *S*_*j*_, *j* = 1 … *n*, and *P*_*i*_, *i* = 1 … *m*, are the concentrations of the participating substrates and products. The reaction’s equilibrium constant, *K*_*eq*_, represents the ratio of the concentrations of substrates, 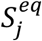, and products,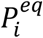, at the equilibrium, and it can be expressed as:

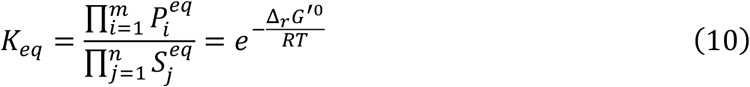

where Δ_*r*_*G*^′0^ denotes the standard Gibbs free energy of the reaction, *R* is the universal gas constant and *T* is the temperature. The Gibbs free energy of the reaction can be expressed as

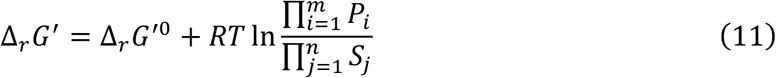

Combining Eqs 9-11, Γ can be expressed as:

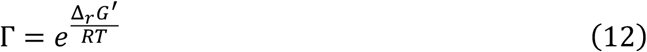

Therefore, for reactions operating towards production of products 𝑃_*i*_, the Gibbs free energy difference is negative and Γ can take values from the interval [0, 1], whereas for reactions operating towards production of substrates *S*_*j*_, Δ_*r*_*G*^′^ is positive and Γ ∈ [1, +∞]. Reactions with values of Γ close to 1 are operating near thermodynamic equilibrium, whereas reactions with Γ ≈ 0 and Γ → +∞ are operating strictly in the forward and backward direction, respectively.

The displacement Γ of a reaction can be determined by knowledge of the Δ_*r*_*G*^′0^,and equivalently the *K*_*eq*_, and the concentrations of the metabolites (substrates and products). The Gibbs free energy of reactions (Δ_*r*_*G*^′0^) can be determined by experiments^*38*^ or it can be estimated using estimation methods, such as group contribution methods.^*39, 40*^

### 2.3. Mass Balances of Enzyme Complexes in Enzymatic Reactions

For a generic multisubstrate multi-product enzymatic reaction

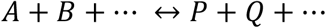

the mass balance equations of the concentrations of enzyme states have the following form:

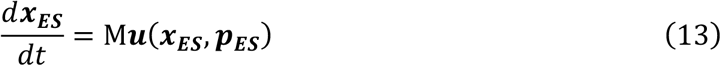

where 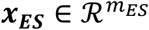 denotes the vector of enzyme states’ concentrations, 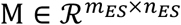 is the stoichiometric matrix describing the dependency of ***x***_***ES***_ and the fluxes of elementary reactions steps 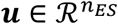, and 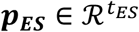 is the parameter vector

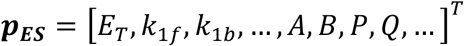

that contains the conserved concentrations of total enzyme, *E*_*T*_, the rate constants of elementary reaction steps, *k*_1*f*_, *k*_1*b*_, …, and other parameters such as the concentrations of substrates and products, *A*, *B*, *P*, *Q*, ….

We assume that an enzyme is not consumed nor produced in the course of a reaction, i.e., the total amount of enzyme, *E*_*T*_, remains constant, i.e., *E* + *EA* + *EB* + ··· + *EP* + ··· = *E*_*T*_. As a consequence, M is a rank deficient matrix as the concentration of an enzyme state can be expressed as a function of other enzyme states. Therefore, the vector of enzyme states, ***x***_***ES***_, can be split in the dependent enzyme states, ***x***_***ES,d***_, and the independent enzyme states, ***x***_***ES,i***_, and the temporal evolution the enzyme states can then be expressed as

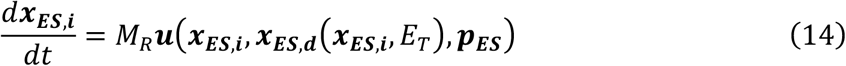

where *M*_*R*_ is a reduced stoichiometric matrix.^*26, 41*^

### 2.4. Sampling of Enzyme States and Displacements of Elementary Reaction Steps from Thermodynamic Equilibrium

For exploring both enzyme states’ space and thermodynamic displacements space, the samples can be drawn from a n-variate Dirichlet distribution. A n-variate Dirichlet distribution with all parameters equal to one allows generating a population of uniformly distributed samples over n-dimensional simplices.^*19*^

#### 2.4.1. Sampling of Enzyme States

In the space formed by the concentration values of the enzyme in its free form, *E*, and in the form of enzyme-metabolite complexes, *ES*_*i*_ the conservation of *E*_*T*_:

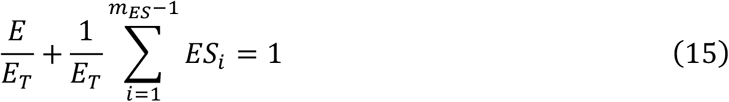

represents a *m*_*ES*_-dimensional simplex. Therefore, the samples of enzyme states scaled by *E*_*T*_ can be efficiently generated by drawing samples from the Dirichlet distribution.^*19*^

#### 2.4.2. Sampling of Thermodynamic Displacements of Elementary Reaction Steps

The overall displacement of a reaction from its equilibrium, Γ, can be expressed as a product of thermodynamic displacements of elementary reactions steps, *γ*_*K*_, belonging to a set *D*:

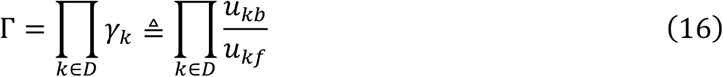

where *u*_*kb*_ is the backward, and *u*_*kf*_ the forward rate of the k^th^ elementary reaction step. The content of the set *D* depends on the kinetic mechanism of a reaction. For example, in the case of ping-pong or ordered kinetic mechanisms, the set *D* contain all elementary steps. From the above expression, we obtain by applying logarithm:

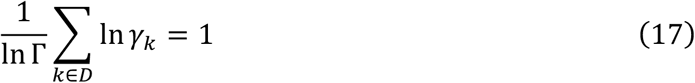

In the space of logarithms of thermodynamic displacements of elementary reactions steps, *γ*_*k*_, Eq. 17 represents a k-dimensional simplex and the samples of *γ*_*k*_ can promptly be generated using the Dirichlet distribution. Observe that for Γ = 1, the thermodynamic displacements of elementary steps are equal to one, i.e., *γ*_*k*_ = 1, *k* ∈ *D*.

Information about the marginal distributions of the enzyme states or displacements from thermodynamic equilibrium of elementary reaction steps inferred from experimental observations can be used to generate refined sets of these quantities. We can accomplish this by adjusting the parameters of the Dirichlet distribution to match the experimental data.^*19*^

### 2.5. Elasticities

Elasticities are defined within the MCA formalism^*9, 10, 12*^ as the local sensitivities of reaction rates to the changes in metabolite concentrations and parameters, and they are required to compute the flux and concentration control coefficients.^*9, 10, 12, 33–35*^

According to the MCA formalism, the elasticities of the enzyme states, ***x***_***ES,i***_, and of the elementary fluxes, ***u***, with respect to the parameters, ***p***_***ES***_, can be expressed in the following form:^*19*^

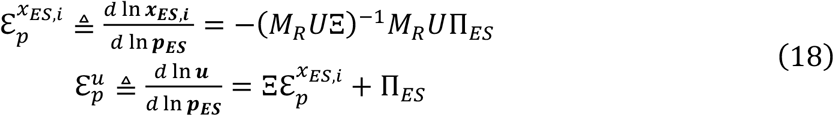

where 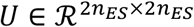 is a diagonal matrix of the forward and backward elementary rates, and 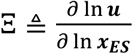 and 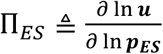 are the matrices of the sensitivities of elementary reaction rates with respect to the enzyme states and the system parameters, respectively. Considering the conservation of the total amount of enzymes, we can write

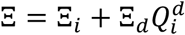

where 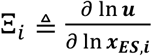 and 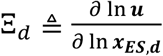 are the matrices of the sensitivities of elementary reaction rates with respect to independent and dependent enzyme states, and 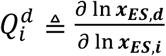 is the relative abundance of the dependent enzyme states with respect to the independent ones.^*19, 33*^

## 3. RESULTS AND DISCUSSION

### 3.1. Analytical Expressions for Elasticities of Several Enzymatic Mechanisms

#### 3.1.1. Ping Pong Bi Bi mechanism

The ping pong mechanism, also called double displacement reaction^*21*^, is characterized by the existence of a substituted enzyme intermediate, *E*^*^, that is temporary formed after the binding of the first substrate, *A*, to the enzyme, *E* (Figure 2b). In the ping pong Bi Bi mechanism, the occurrence of ternary complexes is structurally impossible due to an overlap of the binding sites for the substrates *A* and *B*, and the first product, *P*, is created and released before the second substrate, *B*, binds.^*21*^ Examples of reactions that exhibit the ping pong mechanism include pyruvate carboxylase and serine proteases such as trypsin and chymotrypsin.

**Figure 2.**
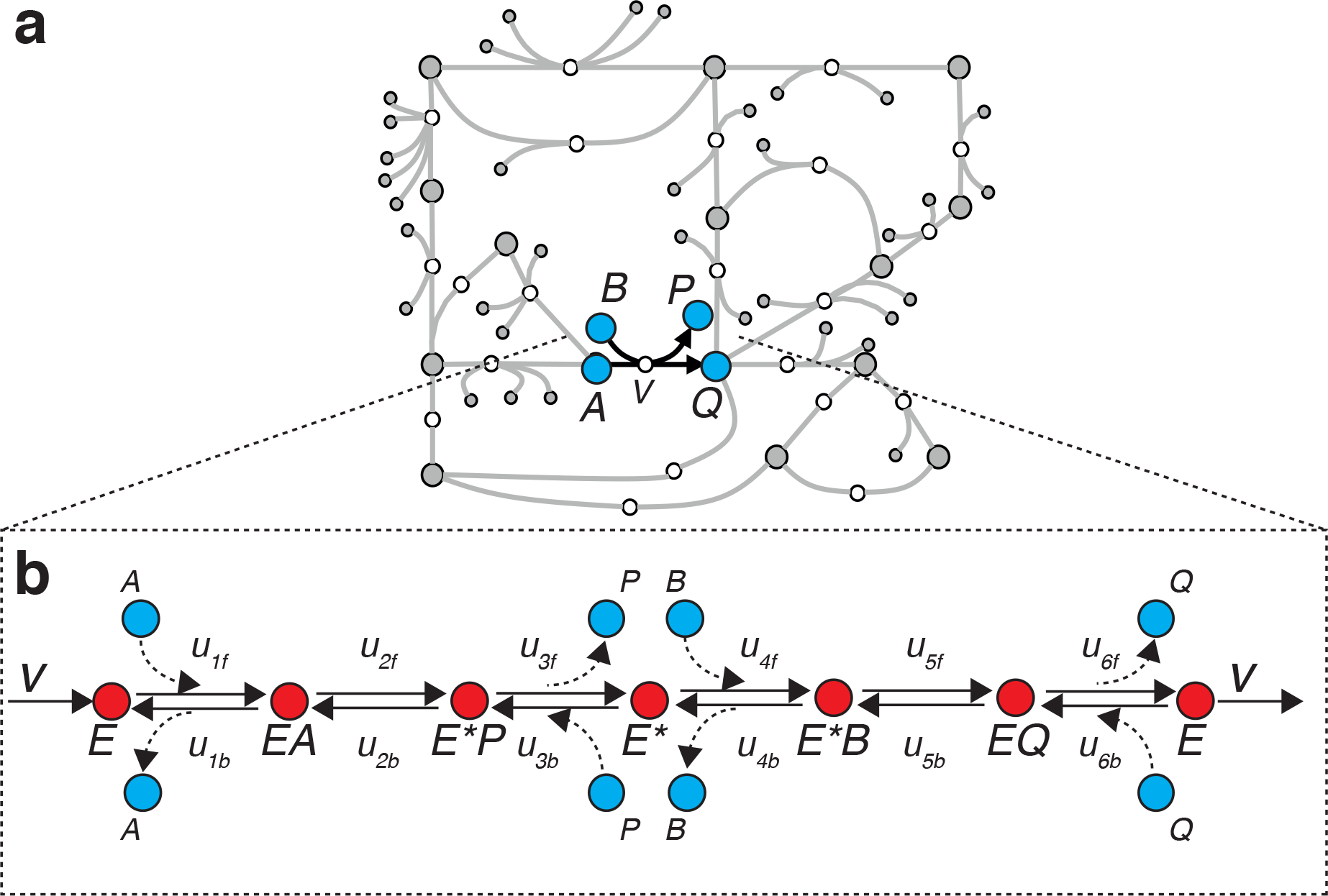
Two representations of a ping pong Bi Bi enzymatic reaction. a) network representation characterized by the flux through the reaction and the metabolite concentrations of participating metabolites; b) more detailed mechanistic representation additionally characterized by the enzymatic mechanism, the enzyme states, and the fluxes of elementary reaction steps.

The net fluxes of six elementary reactions steps, 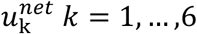, of the ping pong mechanism are all equal to the net flux of the overall reaction, *V*. Therefore, using the definition of the thermodynamic displacement from equilibrium of elementary reaction steps from Eq. 16, the forward and backward elementary reaction rates can be expressed as:

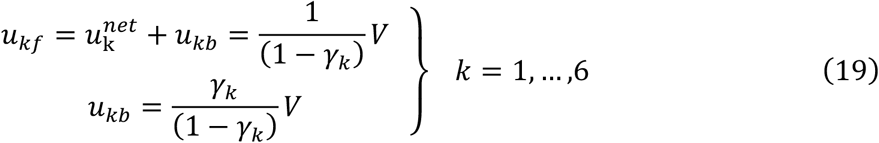

From the conservation of the total amount of enzyme

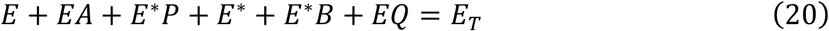

we can consider the enzyme states *EA*, *E*\**P*, *E*\*, *E*\**B*, and *EQ* to be independent and *E* to be dependent.

The elasticities of the ping-pong mechanism can be expressed analytically in the form of Eq. 18, where the stoichiometric matrix of the mechanism is

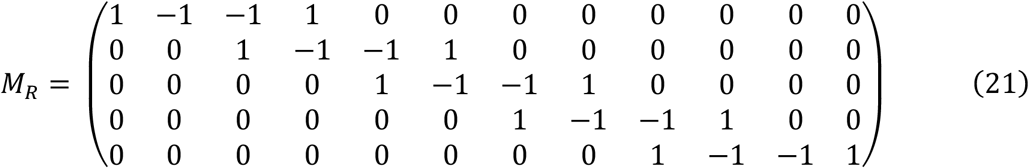

and the matrix of the elementary rates can be derived from Eq. 19:

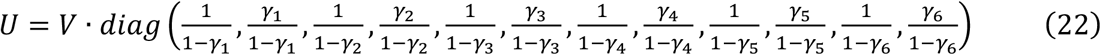

The sensitivities of the elementary reaction rates with respect to the vectors of independent enzyme states [*EA*, *E*\**P*, *E*\*, *E*\**B*, *EQ*] and parameters ***p***_***ES***_ = [*E*_*T*_, *A*, *B*, *P*, *Q*]^*T*^ read as

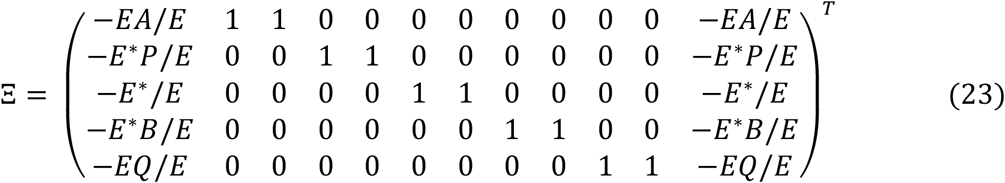

and

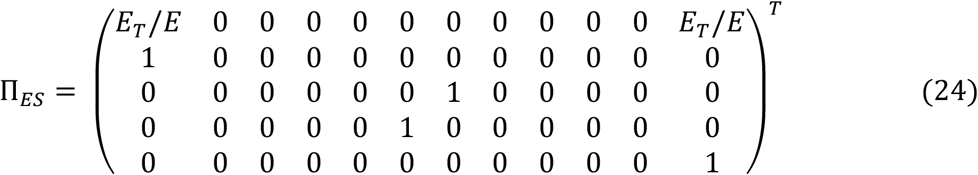

Substituting Eqs. 21–24 into Eq. 18, and considering that

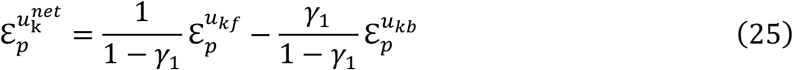

we finally obtain

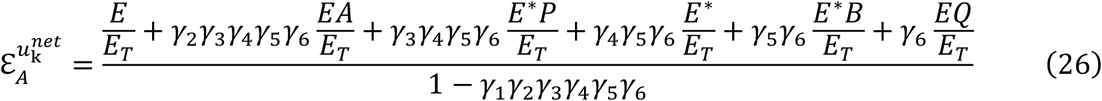

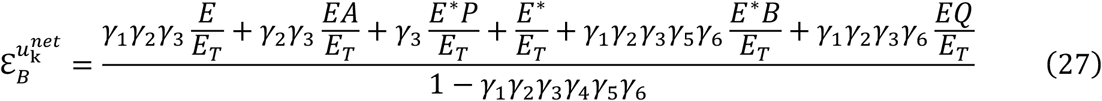

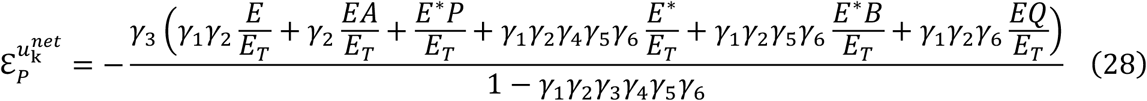

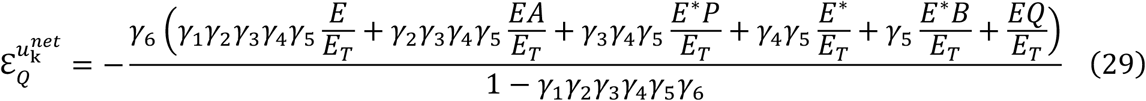

and, as expected, 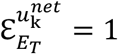.

#### 3.1.2. Ordered Bi Bi mechanism

The ordered Bi Bi mechanism is characterized by a compulsory order of substrate binding and product release (Figure 3). The enzyme *E* first binds to substrate *A*, and creation of the *EA* complex results in the formation of a binding site for substrate *B*. The ternary *EAB* complex is then formed and isomerized to the *EPQ* complex which releases first the product *P* and then the product *Q*. Examples of reactions that follow the ordered bi-bi mechanism include lactate dehydrogenase^*42*^, alcohol dehydrogenase^*43*^, and many other dehydrogenases.

**Figure 3.**
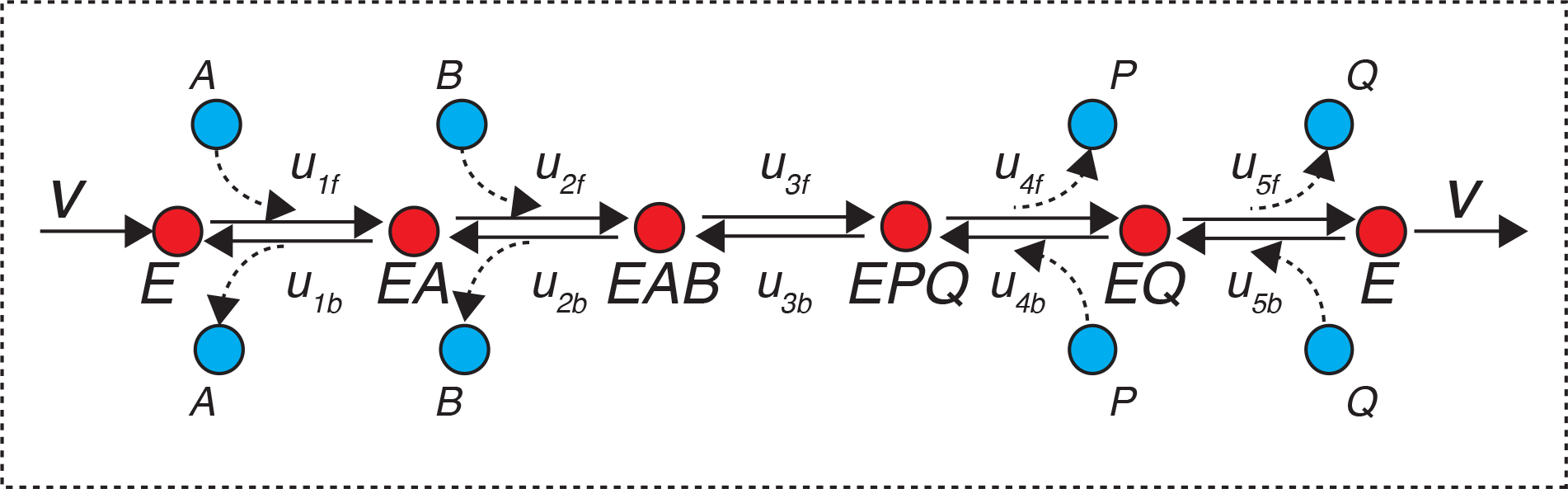
Ordered Bi Bi enzymatic mechanism.

Using the formalism presented in Section 3.1.1., one can obtain the following expressions for the elasticities of the net rates with respect to the parameter vector ***p***_***ES***_ = [*A,B,P,Q*]^*T*^:

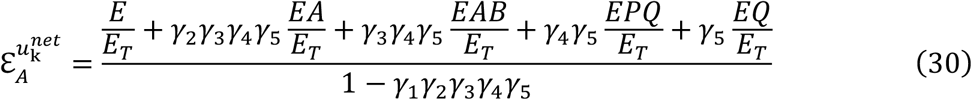

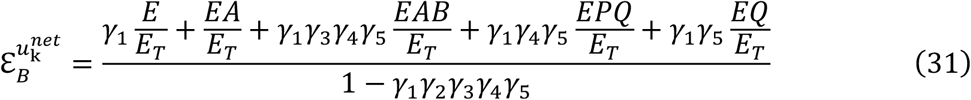

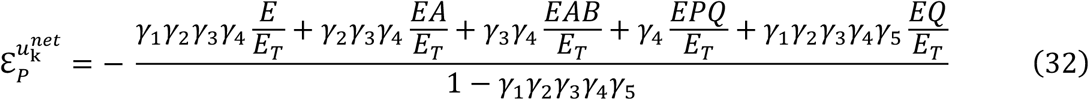

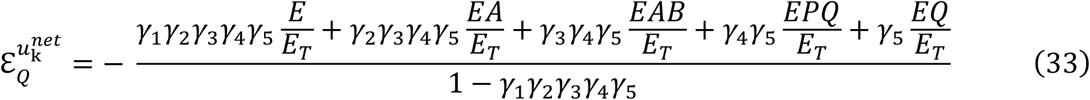

#### 3.1.3. Ordered Uni Bi mechanism

The ordered Uni Bi mechanism is characterized by a compulsory order of product release (Figure 4). Examples of reactions that are conform with the ordered uni-bi mechanism include fructose-bisphosphate aldolase^*44*, *45*^ and isocitrate lyase^*46*^.

**Figure 4.**
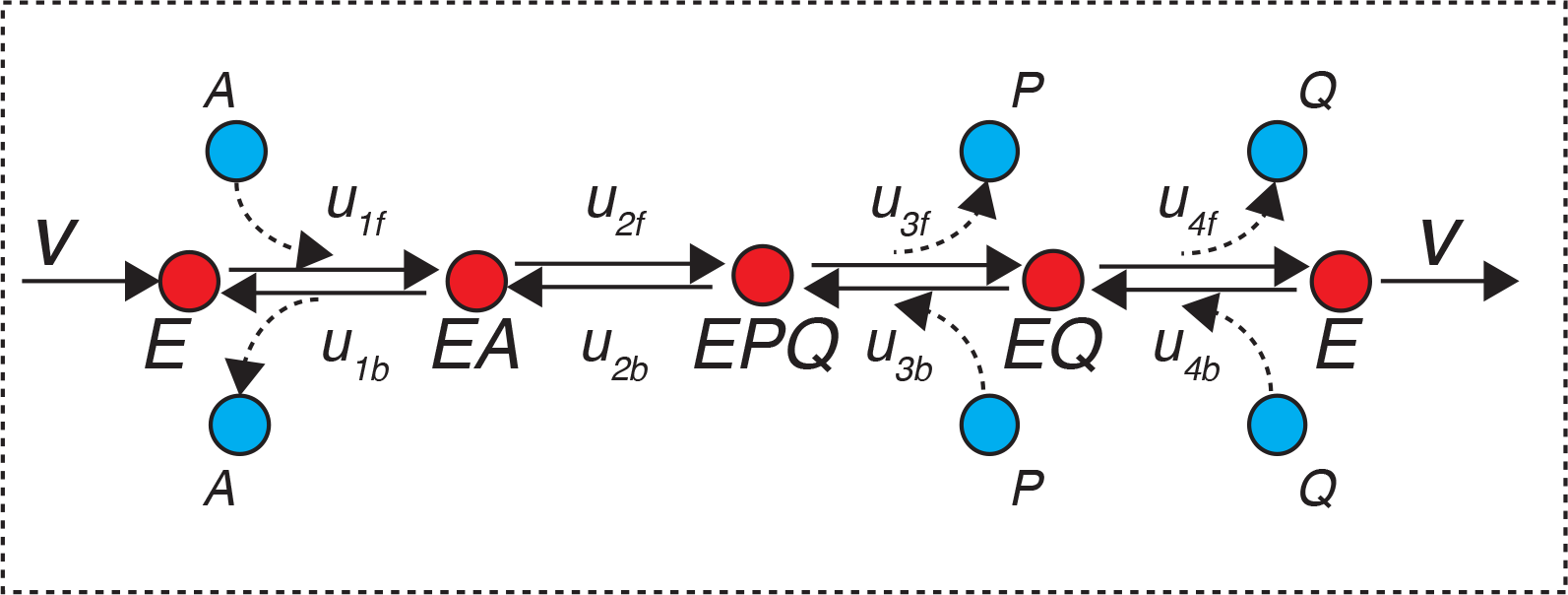
Ordered Uni Bi enzymatic mechanism.

The elasticities of the net rates with respect to the parameter vector ***p***_***ES***_ = [*A,P,Q*]^*T*^ for this mechanism read as:

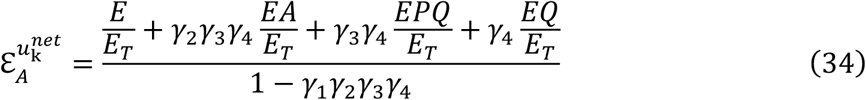

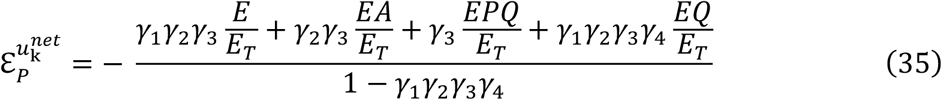

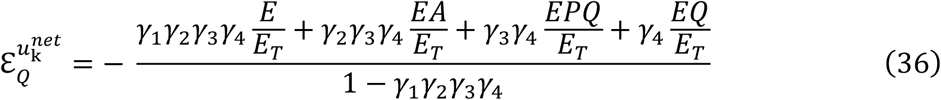

#### 3.1.4. Ordered Bi Uni mechanism

The ordered Uni Bi mechanism is characterized by a compulsory order of substrate binding (Figure 5). Examples of reactions that exhibit this mechanism include 3-methylaspartate ammonia-lyase^*47*^ and pyruvate aldolase^*48*^.

**Figure 5.**
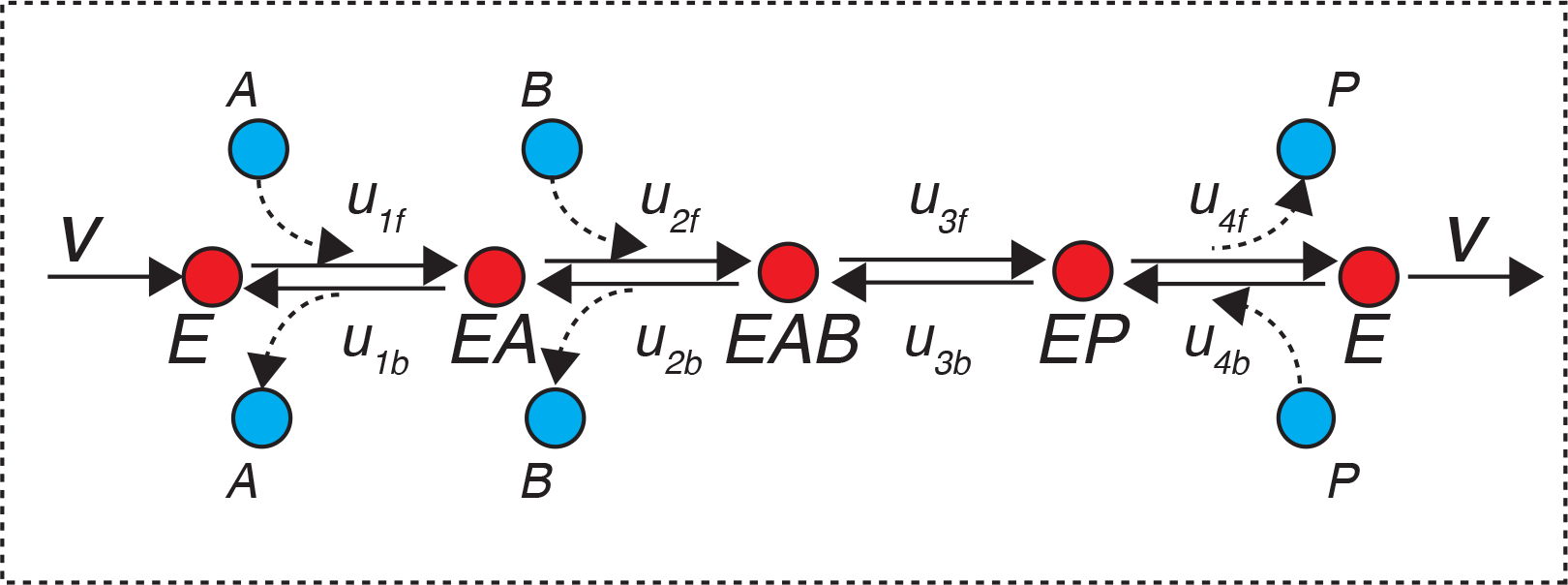
Ordered Bi Uni enzymatic mechanism.

The elasticities of the net rates with respect to the parameter vector ***p***_***ES***_ = [*A*, *B*, *P*]^*T*^ for this mechanism can be written as:

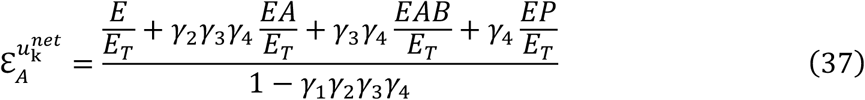

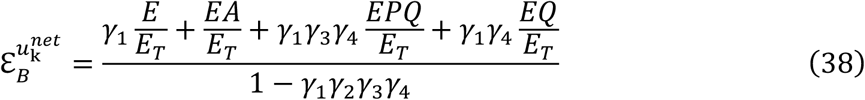

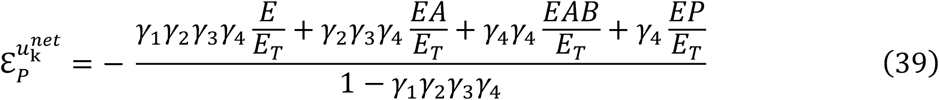

### 3.2. Schematic Method for Deriving Analytical Expressions for Elasticities

The algebraic manipulations used in deriving the analytical expressions for elasticities (Section 3.1) are rather complicated for mechanisms with a large number of enzyme states. However, inspection of the analytical expressions from Section 3.1. reveals a regularity in terms that multiply the enzyme states thus providing a way to shorten the derivations substantially. We propose here a schematic method that is reminiscent of Mason’s gain formula^*22*^ for finding the transfer function of linear systems from the control theory. We present the method through an elementary example.

#### 3.2.1. Illustrative example

Consider the reversible Uni Uni mechanism

**Figure 6.**
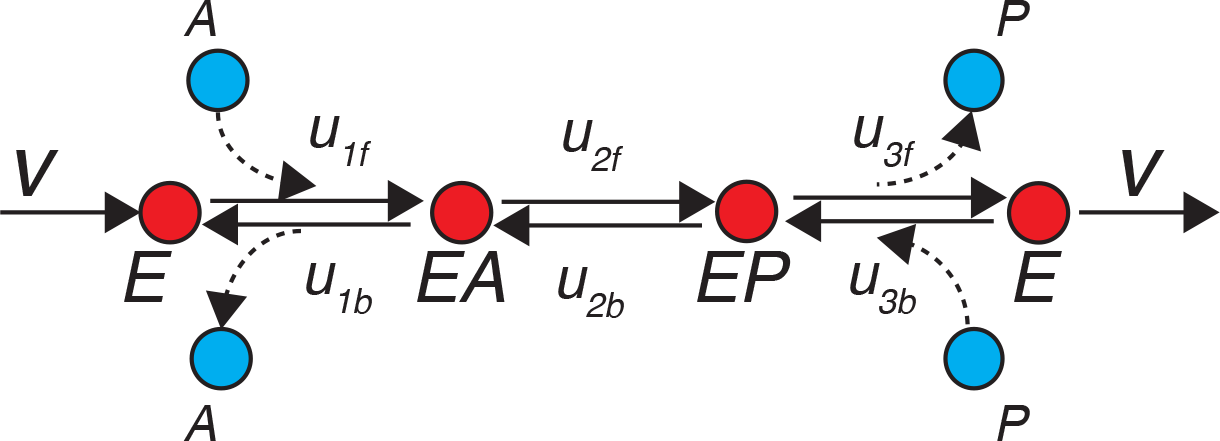
Reversible Uni Uni mechanism

The analytical expressions for the elasticities of the net rates of this mechanism with respect to the parameters ***p***_***ES***_ = [*A*, *B*]^*T*^ are derived elsewhere:^*19*^

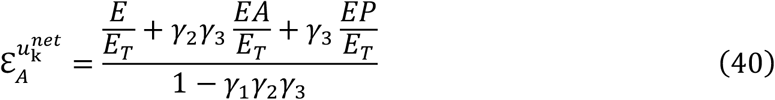

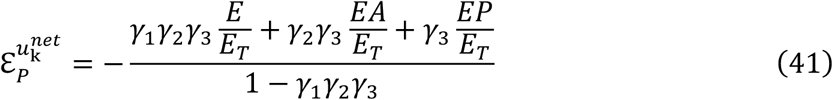

The expressions Eqs. 40-41 can be obtained schematically in two steps as follows.

<u>Step 1</u>: Draw a schematic diagram of the kinetic mechanism as a graph, where the enzyme states that both (i) bind with a substrate and (ii) are formed by releasing a product are represented by two nodes. Such a state in the illustrative example is the enzyme in its free form (Figure 6). We distinguish these two nodes as the input node, that connects to a substrate (Figure 7a, empty circle) and the output node that connects to a product (Figure 7a, full circle). The other enzyme states, *EA*/*E*_*T*_ and *EP*/*E*_*T*_, are represented with a node of the graph (Figure 7a).

**Figure 7.**
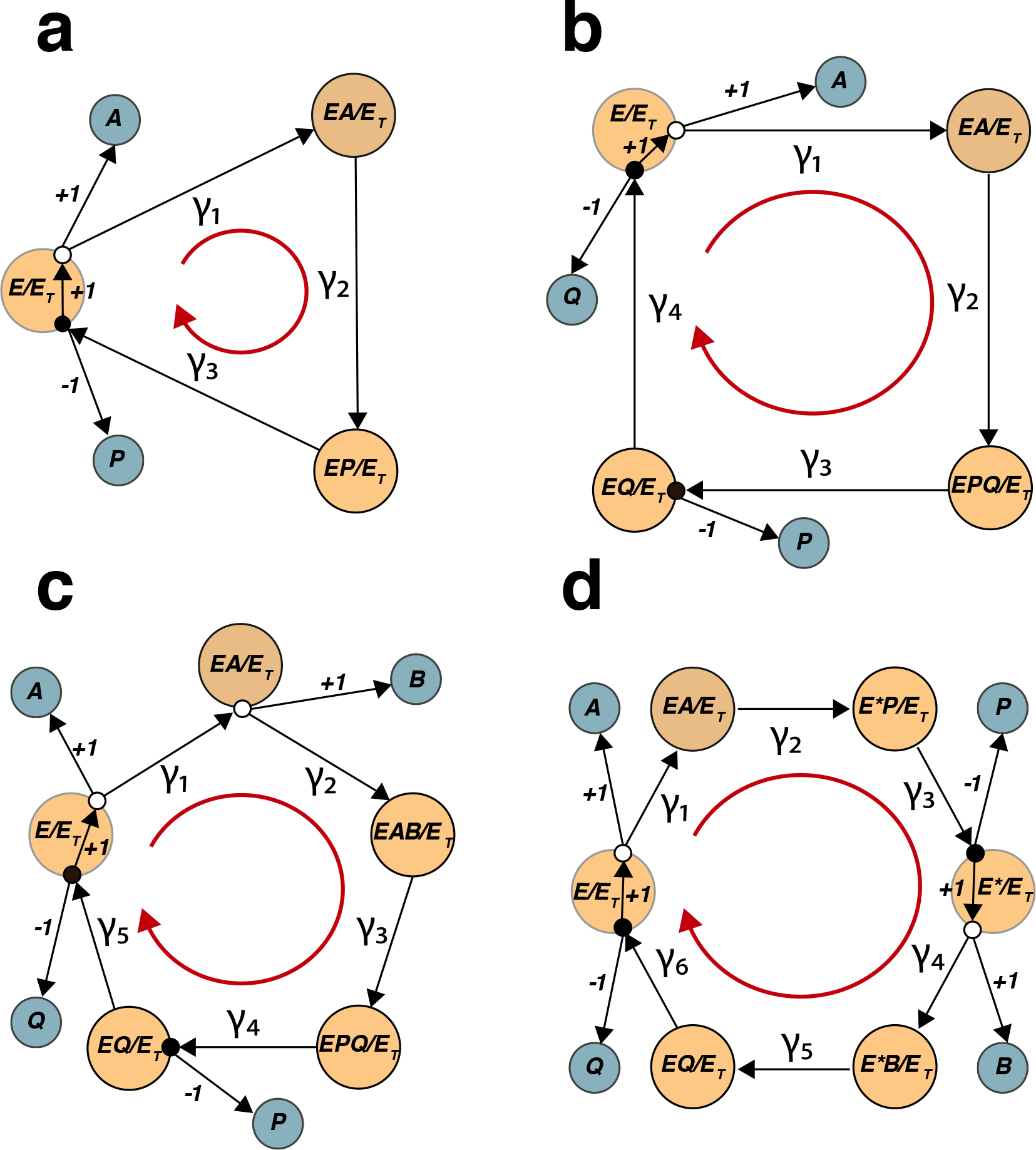
Directed graphs of typical enzymatic mechanisms: Uni Uni mechanism (a), Ordered Uni Bi Mechanism (b), Ordered Bi Bi mechanism (c), and Ping Pong Bi Bi mechanism (d).

It is assumed that there is a net production of the product *P*, i.e., a net flow is directed from *A* to *P* (Figure 7a). The vertices of the graph that connect each two enzyme states are weighted by the equilibrium displacements of the elementary steps connecting them, and they are directed along the assumed net flow. For example, the vertex that connects *E*/*E*_*T*_ and *EA*/*E*_*T*_ has a weight of γ_1_, and it is directed from *E*/*E*_*T*_ to *EA*/*E*_*T*_. The vertices that connect a substrate to the input node of an enzyme state have the weight of +1, whereas the vertices that connect the output nodes of an enzyme state to a product are directed toward the product and have the weight of −1. The vertices that connect the input and the output node of an enzyme state are directed towards the input node and have the weight of +1.

<u>Step 2</u>: From the directed graph formed in Step 1 (Figure 7a), the analytic expression of the elasticity of the net rate with respect to *A* and *P*,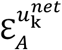 and 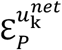, can be obtained as follows.

The denominator of 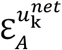 and 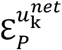 is equal, and we form it by subtracting from 1 the product of the weights of the vertices that form the graph loop, i.e., we obtain 1 – *γ*_1_*γ*_2_*γ*_3_ from Figure 7a.

To derive the terms in the numerator, we start by forming for each of enzyme states the direct path from that state to *A* when we derive 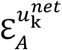 or to *P* for 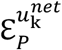. For the enzyme states with two nodes, the direct path starts from the input node. Then, to form the numerator terms, the weights of the direct paths are multiplied with the corresponding enzyme state. The summary of the procedure for the example is provided in Table 1.

**Table 1.** Direct paths of the directed graph used to form the numerators of 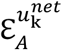 and 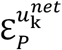 of the reversible Uni Uni mechanism.

|  | Direct path |  | Gain | Corresponding numerator term |
| --- | --- | --- | --- | --- |
|  | From | To |  |  |
| $\mathcal{E}_A^{u_k^{net}}$ | $E/E_T$ | $A$ | 1 | $E/E_T$ |
| | $EA/E_T$ | $A$ | $\gamma_2\gamma_3$ | $\gamma_2\gamma_3EA/E_T$ |
| | $EP/E_T$ | $A$ | $\gamma_3$ | $\gamma_3EP/E_T$ |
| $\mathcal{E}_P^{u_k^{net}}$ | $E/E_T$ | $P$ | $-\gamma_1\gamma_2\gamma_3$ | $-\gamma_1\gamma_2\gamma_3E/E_T$ |
| | $EA/E_T$ | $P$ | $-\gamma_2\gamma_3$ | $-\gamma_2\gamma_3EA/E_T$ |
| | $EP/E_T$ | $P$ | $-\gamma_3$ | $-\gamma_3EP/E_T$ |

#### 3.2.2. Schematic Method

The method can be summarized as follows. Create a directed graph representing a kinetic mechanism as described in Step 1 of Section 3.2.1. The analytical expression for the elasticities of the reaction rate of the mechanism with respect to its substrates and products can be readily derived from the expression

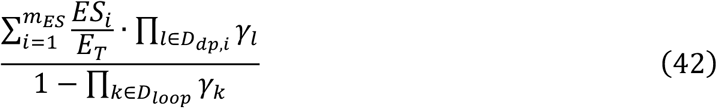

where *D*_*loop*_ is the set that contains the vertices that form the graph loop, *D*_*dp,i*_ the set of vertices that form the direct path from the enzyme state *ES*_*i*_ towards the metabolite of interest.

#### 3.2.3. Ping Pong Bi Bi Mechanism

An interesting example for the application of the schematic method is the ping pong bi bi mechanism because it has two enzyme states, the free enzyme, *E*, and the substituted enzyme intermediate, *E*^∗^, that have two nodes (Figure 7d).

From the directed graph (Figure 7d) and Eq. 42, we obtain for the denominator 1 – *γ*_1_*γ*_2_ *γ*_3_*γ*_4_ *γ*_5_*γ*_6_, and the terms of the numerators are provided in Table 2.

**Table 2.**
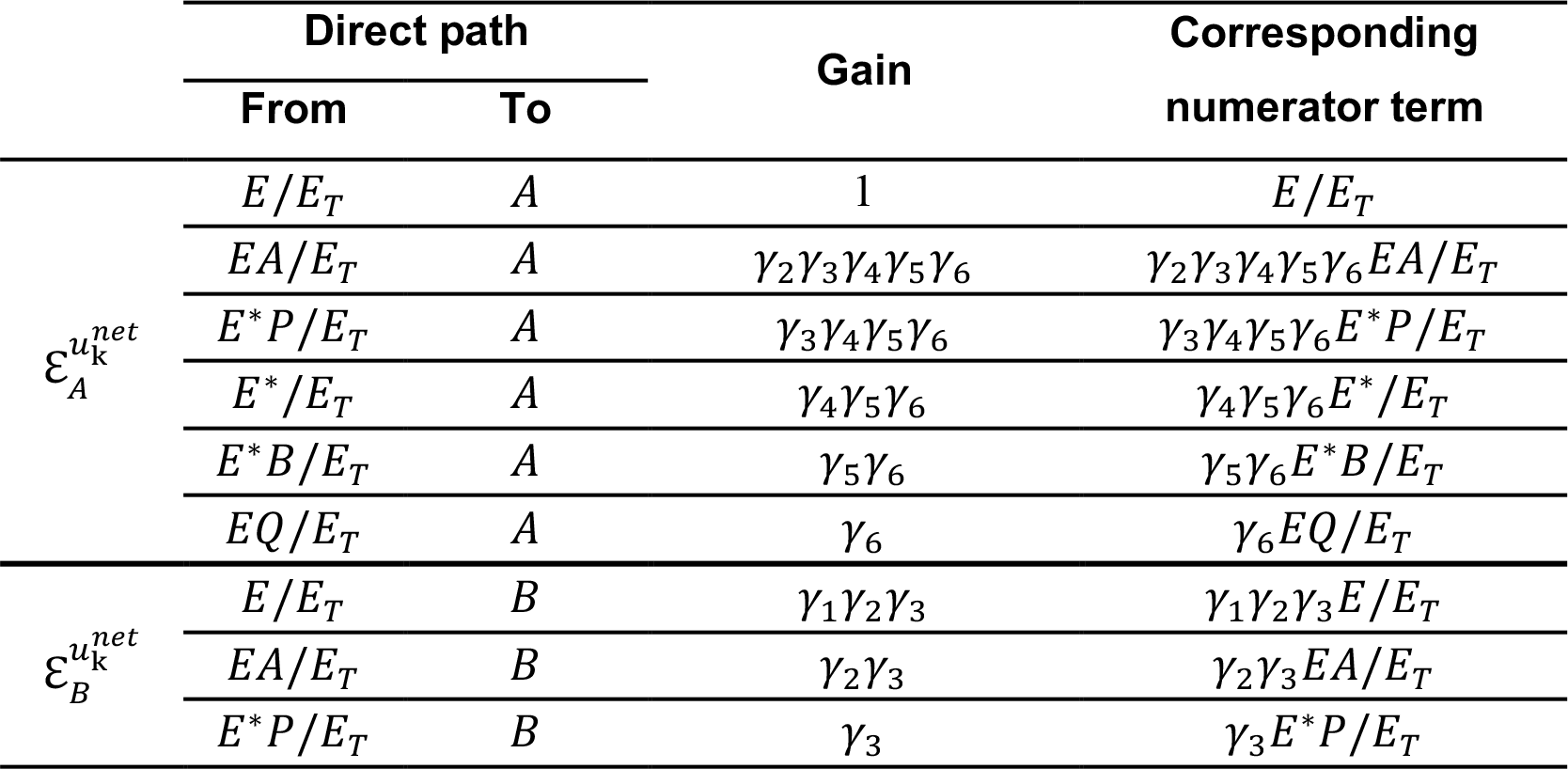
Direct paths of the directed graph used to form the numerators of 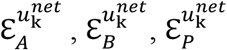 and 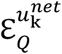 of the ping pong Bi Bi mechanism.

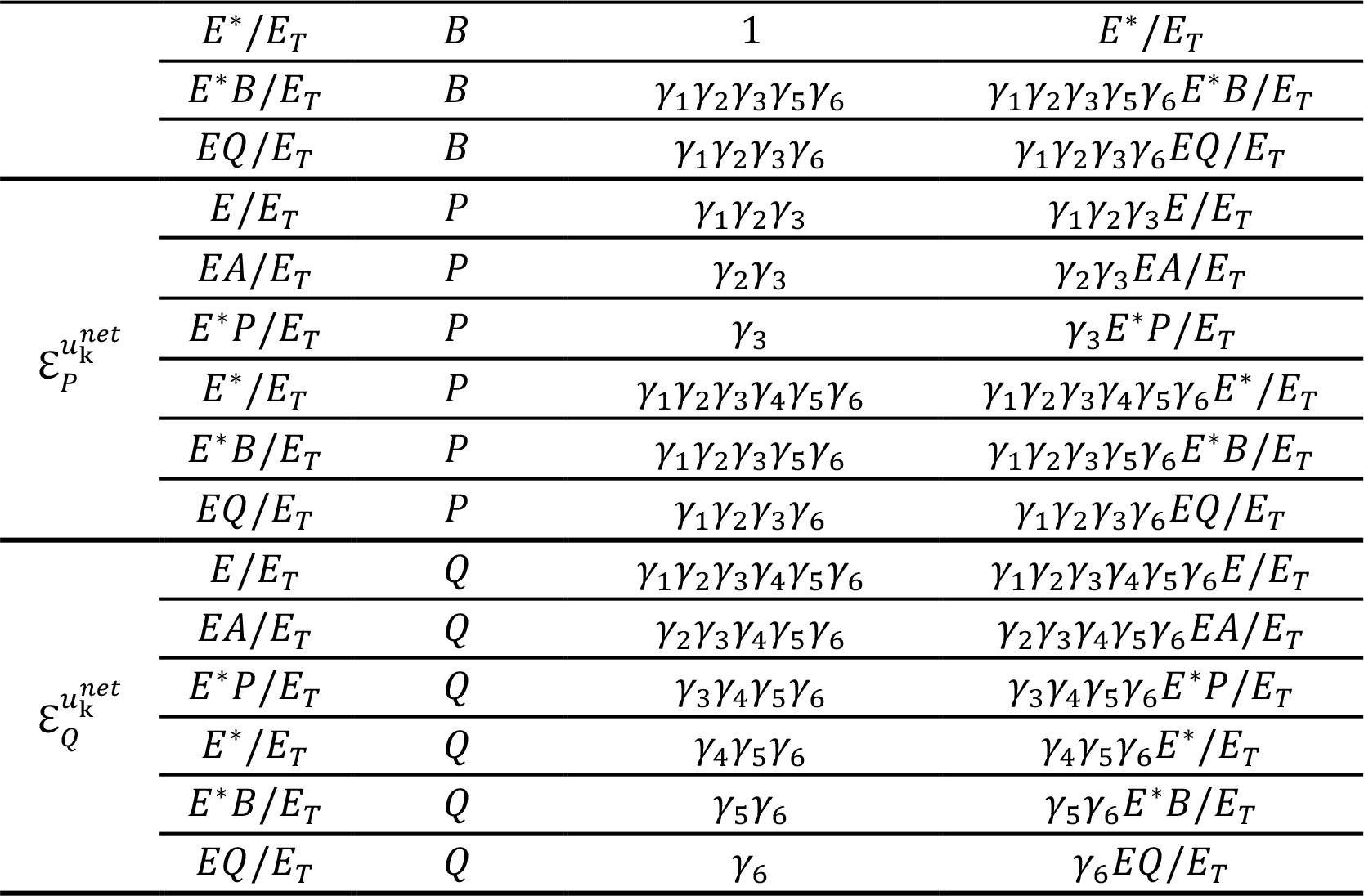

In a similar way, one can derive the analytic expressions for the elasticities of the ordered Uni Bi and ordered Bi Bi mechanisms using the directed graphs from Figure 7b and 7c, respectively (Supplementary material).

## 4. CONCLUSIONS

The uncertainties and absence of data about kinetic properties of enzymes remain a major hurdle for developing kinetic models of biochemical networks. The construction of these models requires approaches that consider networks as a whole and also consider the mechanistic properties of enzymes. Indeed, the network quantities such as metabolic fluxes and metabolite concentrations affect the behavior of elementary reaction steps within kinetic mechanisms, and conversely, the parameters and variables corresponding to individual kinetic mechanisms affect the overall behavior of biochemical networks. The formalism based on Monte Carlo sampling of enzyme states and equilibrium displacements of elementary reaction steps discussed here allows us to model the kinetic responses of enzymes despite the lack of kinetic data, and therefore, to model the effects of individual kinetic mechanisms on metabolic networks. The formalism, coupled together with the network information obtained from methods such as TFA, allow us to predict the responses of biochemical networks to genetic and environmental variations. Efficient sampling procedures for generating missing kinetic data used in the formalism represent a valuable tool for methods that use Monte Carlo sampling to generate populations of large-scale kinetic models^*36, 37, 49–62*^.

Though the proposed schematic method for deriving the analytical expressions for elasticities can cover a wide gamut of the ordered and ping-pong mechanisms, its extension to the random-order and other more complex mechanisms remains to be addressed.

## ASSOCIATED CONTENT

### Supporting Information

The following files are available free of charge.

Further illustration of the schematic method for derivation of the elasticities for: (i) ordered Uni Bi; (ii) ordered Bi Uni and (iii) ordered Bi Bi mechanisms (DOCX)

## AUTHOR INFORMATION

### Author Contributions

The manuscript was written through contributions of all authors. All authors have given approval to the final version of the manuscript.

### Notes

The authors declare no competing financial interests.

### ABBREVIATIONS

MCA: metabolic control analysis
TFA: thermodynamic-based flux analysis
EBA: energy balance analysis
NET: network-embedded thermodynamic analysis

